# Detection of mismatching visual kinematics during visuomotor incongruence: A comparison of spatially matched delays vs offsets along identical trajectories

**DOI:** 10.1101/2025.06.18.660374

**Authors:** Fanni Peters, Peng Wang, Jakub Limanowski

**Affiliations:** Institute of Psychology, University of Greifswald, Greifswald, Germany

## Abstract

Introducing delays to visual movement feedback or displacing it in space is a common experimental manipulation to study the neurocomputational basis of self-other distinction. While both manipulations imply spatial (positional) mismatches, there are crucial differences between them, such as the asynchrony of higher-level visual vs motor kinematics (velocity, acceleration etc.) resulting from added time delays. In this preregistered experiment, we used a continuous elliptical drawing task while presenting visual movement feedback that was congruent or incongruent; i.e., with an added constant time delay or offset. The task allowed the presentation of delayed and offset feedback along identical trajectories, while controlling for the presence of a spatial discrepancy by matching delay and offset levels. Detection and discrimination of delays and offsets was remarkably similar during execution, suggesting comparable perceptual thresholds for visuomotor mismatches resulting from asynchronous vs nonbiological visual kinematics. During mere passive observation of the same feedback, offsets could also be detected above chance level, in line with the known visual sensitivity to violations of kinematic invariants. However, the detection of offsets at medium levels was better during execution, suggesting a benefit of action for identifying nonbiological kinematics in visual movement feedback.

## Introduction

Distinguishing sensations that have been caused by one’s actions from those that have been caused by others is a key prerequisite for agency and control (Tsakiris et al., 2005; Synofzik et al., 2008; Salomon et al., 2013; Haggard, 2017; Wen et al., 2018; Wen & Imamizu, 2022; Limanowski, 2022). The brain likely achieves this by calculating a prediction of the sensory consequences of (planned) actions by means of a “forward” model, and comparing these predictions with the actual sensory reafference (Miall et al., 1996; Wolpert & Miall, 1996): Sensory feedback matching the predictions is perceived as a result of one’s own actions. In contrast, the registration of unpredicted sensations is thought to lead to a loss of perceived control or agency, and the attribution of movement generation to someone else (Frith et al., 2000; Farrer & Frith, 2002; Tsakiris & Haggard, 2005).

To understand the underlying processes and their neurocomputational foundations, researchers have artificially distorted sensory movement feedback, for instance, by displacing the seen movement in time (i.e., by adding delays) or space (e.g., by adding angular deviations or mirroring). There is an ongoing debate about whether “temporal” and “spatial” visuomotor comparisons provide equally strong cues for the self-or-other attribution of observed movements (cf. Farrer et al., 2008; Krugwasser et al., 2019). This debate centers on the importance of congruent visual and motor kinematics and their relation to action timing: Delaying and displacing visual movement feedback both imply spatial (i.e., positional) mismatches between the moving body part and the visual feedback. However, a key difference is that displacement preserves all visual kinematics in “real-time” (synchronously), merely requiring the spatial remapping of positional coordinates (Rohde & Ernst, 2016; Limanowski, 2026). In contrast, added delays imply asynchronous motor and visual kinematics; e.g., a motor acceleration will produce a visual acceleration only after the time delay (ibd.). Along these lines, it has been argued that artificially introduced feedback delays may provide stronger cues for self-other distinction than spatial manipulations, because they imply changes to action timing rather than merely positional recalibration (Rohde & Ernst, 2016; cf. Tanaka & Imamizu, 2025).

Testing this experimentally, however, is notoriously difficult. While many studies have displaced visual movement feedback in space or time, only few studies have attempted their comparison in a single experiment (see Farrer et al. 2008; Krugwasser et al. 2019). Furthermore, the typical spatial manipulation is angular displacement or mirroring. This implies the visual feedback following a different trajectory than movements—introducing a crucial difference to delayed movement feedback conditions, and limiting the interpretability of the resulting commonalities and differences.

Therefore, in this preregistered study, we used a novel elliptical drawing task that allowed a systematic comparison of perceptual thresholds for spatial (offset) and temporal (delay) incongruities in visual movements, and the influence of action on their detection and discrimination. Participants continually drew ellipses, with congruent or manipulated visual movement feedback. The key novelty of our design was that all movements were executed along an identical, elliptical trajectory, which the visual movement feedback reflected congruently, or followed with a constant time delay (i.e., with a spatial discrepancy and asynchronous kinematics) or a constant spatial offset (i.e., with a spatial discrepancy but synchronous kinematics). We matched delayed vs offset visual feedback, at six levels each, in terms of their average Euclidean distance from the executed movement. Thus, our design allowed us to compare the perceptual sensitivity to delays vs offsets along matching visual movement trajectories; while controlling for the presence of a spatial discrepancy. Furthermore, it allowed us to employ a three-alternative-choice response format; i.e., in contrast to previous work, participants were not just asked to indicate whether the visual movement feedback was congruent or incongruent, but to specifically identify the kind of incongruence (delay or offset).

Delays and offsets along identical trajectories may be indistinguishable if movements are performed with constant velocity (e.g., circular or sinusoidal movements, Rohde & Ernst, 2016). Therefore, to manipulate the correspondence of visual and motor kinematics as the key difference between delayed and offset visual movement feedback, we sought to introduce systematic variations in movement velocity. By using an elliptical trajectory, we capitalized on kinematic invariants in movement production as described by the “two-thirds power law” (Binet & Courtier, 1893); i.e., that biological movements tend to be slower at trajectories with stronger curvature (Box S1; cf. Viviani & Stucchi, 1992; de’Sperati & Stucchi, 1995; Dayan et al., 2007; Hun & Sejnowski, 2015). Crucially, these invariants were preserved in the delayed feedback condition, but shifted along the elliptical trajectory in the offset feedback conditions. As the presence of a spatial discrepancy was controlled for, differences in detection performance could, therefore, specifically be related to the sensitivity to mismatching (asynchronous) visual and motor kinematics.

Although the visual kinematics in our offset condition were synchronous, they contained another visual “error signal” that could be used for visuomotor mismatch detection; namely, nonbiological velocity profiles. Previous work has shown that the visual system is sensitive to violations of kinematic invariants such as those described by the two-thirds power law (Salomon et al., 2016; Fraser et al., 2025; cf. Knoblich & Prinz, 2001; Knoblich et al., 2002). Therefore, we included a visual control condition (participants observing their visual movement playback, with an identical response format) to establish whether action would improve the detection of manipulated visual movements (asynchronous or nonbiological visual kinematics).

Our key hypotheses were (see Table S1 for detailed preregistered hypotheses): Firstly, during execution, delays and offsets should, on average, be detected above chance level; while accuracy should improve with increasing amount of mismatch. During observation, we expected the playback of offsets, but not delays to be recognized above chance level. This followed from the fact that offset movements violated biological kinematic laws (see above), whereas delayed movements, when merely played back, were indistinguishable from congruent movements. Secondly, we expected better detection performance, and a stronger performance increase with increasing amount of mismatch, during execution compared with observation. For delays, this prediction was trivial (see above); for offsets, it assumed a benefit of action for the perception of visuomotor mismatches resulting from nonbiological visual kinematics.

## Methods

Unless explicitly noted otherwise, we preregistered our hypotheses (https://osf.io/39tch) and experimental design and procedure (https://osf.io/udxth) on OSF; we preregistered after having started data acquisition but before inspecting any data or performing any statistical analyses.

### Participants

33 right-handed volunteers (23 women; aged 18–42, mean=24±4.79 years) completed the experiment. All participants provided written informed consent, had normal or corrected-to-normal vision, and reported no history of psychiatric or neurological conditions. The sample size was determined with an a priori power analysis (not preregistered) using G*Power, and based on data from a pilot experiment (*N*=5). This analysis was based on a simplified model using multiple linear regression, as G*Power does not support generalised linear mixed models (GLMM). The target effect corresponded to the difference between the two experimental parts (execution and observation), as quantified in the pilot data. With a medium effect size of Cramér’s V=0.3, and assuming a significance level ⍺ = 0.05 and a target power (1-β) of 0.8, the required sample size was 33. The data of one participant had to be excluded, as no trials in one of the conditions survived our exclusion criteria (see below); resulting in a final sample size of 32. To verify that this sample size was sufficient, we calculated an additional, post-hoc power analysis based on our GLMMs, using the R package “simr” (Green & MacLeod, 2016). For our sample size (N=32) and observed effect sizes (see Results), the estimated power (1-β) was greater than 0.99 in each case (500 Monte Carlo simulations per main and interaction effect, significance level set at 0.05). The experiment was approved by the ethics committee of the University Medicine of Greifswald.

### Apparatus and stimuli

The experimental environment was run on a computer using PsychoPy (version 2024.1.5; Peirce et al., 2019). Participants sat at a table at about 50 cm distance from the computer screen (HP E27 G5 FHD, 27-inch IPS panel, 1920 x 1080 pixels resolution, 75 Hz refresh rate), holding a pen in their right hand with which they had to perform the elliptical movements in the execution phase (see below). The movements were recorded at 75 Hz by a tablet (Wacom, model no: CTL-672, connected to the computer via USB; Fig. 1A). To avoid visual distractions, the tablet and the participant’s right hand were occluded under a cover. Due to the dimensions of the pen, this setup did not allow us to place the monitor horizontally; i.e., directly above the moving hand. Placing the monitor vertically required additional visuomotor coordinate transformations, but in our within-subject design, those were balanced across conditions.

**Figure 1.**
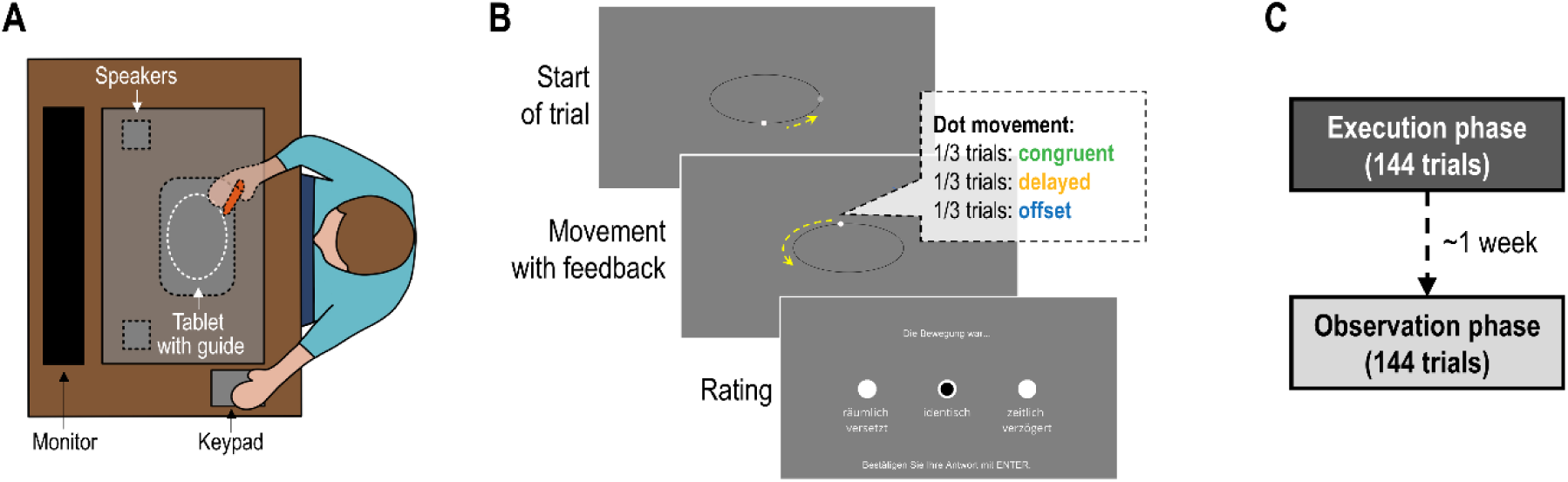
**A:** Experimental setup. Participants sat at a table in front of a computer monitor and performed elliptical pen movements on a graphics tablet, guided by a 3D-printed template, with visual movement feedback provided on the screen. The tablet, template, pen, and the participants’ right hand were occluded from view; to mask any auditory movement feedback, brown noise was played via speakers, also hidden from view. Participants gave rating responses with their left hand via a small key pad. **B:** Schematic trial structure. In the execution phase, the participants started each trial by moving to a predefined region of the ellipse (schematically indicated by the dashed yellow arrow), upon which a white dot appeared. Then they had to perform continuous elliptical movements at ∼0.33 Hz while receiving visual movement feedback via a white dot moving along the visible ellipse on screen. The movement of the dot could be either veridical (congruent with the executed movement, 1/3 of all trials) or manipulated (delayed with respect to the executed movement, 1/3 of all trials; or spatially offset with respect to the executed movement, 1/3 of all trials). There were 6 different levels of delays and offsets, respectively, which were matched in terms of their average distance between pen and dot on the ellipse (see Methods). Participants were instructed to end the trial by left-hand key press as soon as they were ready to rate the visual movement feedback. Importantly, we used a three-alternative-choice response format; i.e., participants had to rate whether each seen movement had been congruent, delayed, or offset with respect to their actual pen movement on the ellipse. **C:** The experiment consisted of an execution phase (as described above) and an observation phase, separated by about one week. In the observation phase, participants viewed a (randomized) playback of the visual movements without moving themselves; they had to rate whether each seen movement had been from a congruent, delayed, or offset movement trial, as in the execution phase.

The elliptical shape of the executed movements was determined by a custom 3D printed template that guided the pen (via a circular inverted T-shape that moved under the template, see Supplementary material). This prevented the pen to be lifted or to deviate to the outside. On screen, participants saw the same elliptical shape as a black outline, and visual movement feedback as a white dot (1.2° visual angle in diameter). The sizes of the physical and visual ellipses were matched in terms of degrees visual angle from the participant’s view. The size of the major axis of the ellipse was 22.5° visual angle, while the size of the minor axis was 10° visual angle. The eccentricity of the ellipse was Σ=0.9 based on the shape used by Bidet-Ildei and colleagues (2008).

To minimise potential auditory movement feedback, resulting e.g. from the pen scratching on the tablet, brown noise was played via speakers (also hidden from view under the cover) throughout the entire experiment. For the rating responses, participants placed their left hand on a keypad (LogiLink USB Numeric Keypad) so that the ring finger was on the “1” key, the middle finger was on the “2” key, the index finger was on the “3” key, and the thumb was on the “Enter” key.

### Experimental design and procedure

Prior to the actual experiment, participants completed a five-part training session (approx. 20 min) to familiarize themselves with the experimental task and conditions. In the first three training sessions, they practiced movements with congruent visual feedback, as well as with a medium delay (240 ms) and a medium offset (8.5°). In the fourth part, participants practiced all trial conditions in a randomized order and became familiar with the rating format. In the final part of the training, the participants practiced moving the pen continuously at the instructed velocity (0.33 Hz).

Following the preregistered protocol (https://osf.io/udxth), the experiment consisted of two phases: a movement (execution) phase and an observation phase. In the first, execution phase, participants had to perform counter-clockwise elliptical right-hand movements with a pen on a tablet (see above). The instructed velocity for movement execution was 3 s per cycle (0.33 Hz), which was practised extensively (see above). We did not include any pacing signals, as any regular signals would have provided cues to compare dot and/or pen positions against—which would have facilitated the detection of visuomotor manipulations. Participants started each trial by crossing a blinking grey “starting point” at the right apex of the ellipse (Fig. 1B). Then, the white dot disappeared for 1500 ms, which was necessary to buffer the visual movement feedback in the delayed conditions.

During the subsequent movement execution, participants received three different kinds of visual movement feedback: congruent, delayed, or offset. In the congruent visual feedback condition, the movement of the dot on screen displayed the participants’ pen movement in “real-time” (bar intrinsic delays of our setup). In the delayed feedback condition, a time delay was introduced between the executed pen movements and the movements displayed via the dot, by buffering the recorded pen movements and replaying them after a constant time interval. Thus, the dot motion lagged behind the executed movements and did not immediately reflect their higher order kinematics, such as changes in velocity or acceleration. There were six possible levels of delay (13.3, 80, 160, 240, 320, and 400 ms) that we chose following previous related works (Leube et al., 2003; Farrer et al., 2008; Limanowski et al., 2017; Krugwasser et al., 2019).

One of our key novelties was the introduction of visual movement feedback that was spatially displaced (offset) along the same, elliptical movement trajectory. I.e., in the offset feedback condition, a constant spatial discrepancy between the pen movement and the visual feedback was implemented; i.e., the dot followed the pen’s trajectory on the ellipse with a constant Euclidean distance. To do this, for each recorded pen position, we determined the spatially displaced coordinates located behind the pen position on the elliptical trajectory (i.e., displaced clockwise) at a fixed Euclidean distance, using a motion vector-based approach (see Supplementary material). To implement a constant spatial offset following the participants’ elliptical movement, we had to restrict the visual movement to the ideal elliptical trajectory. The visual feedback was not delayed in this condition; i.e., despite a positional mismatch, the visual kinematics were preserved. In other words, the dot’s movements reflected changes in the executed movement kinematics in “real-time” (i.e., as in the congruent condition). This implied a shift of the visual velocity profiles along the trajectory of the ellipse, which violated the Two-thirds power law (see Fig. 2 for a display).

**Figure 2.**
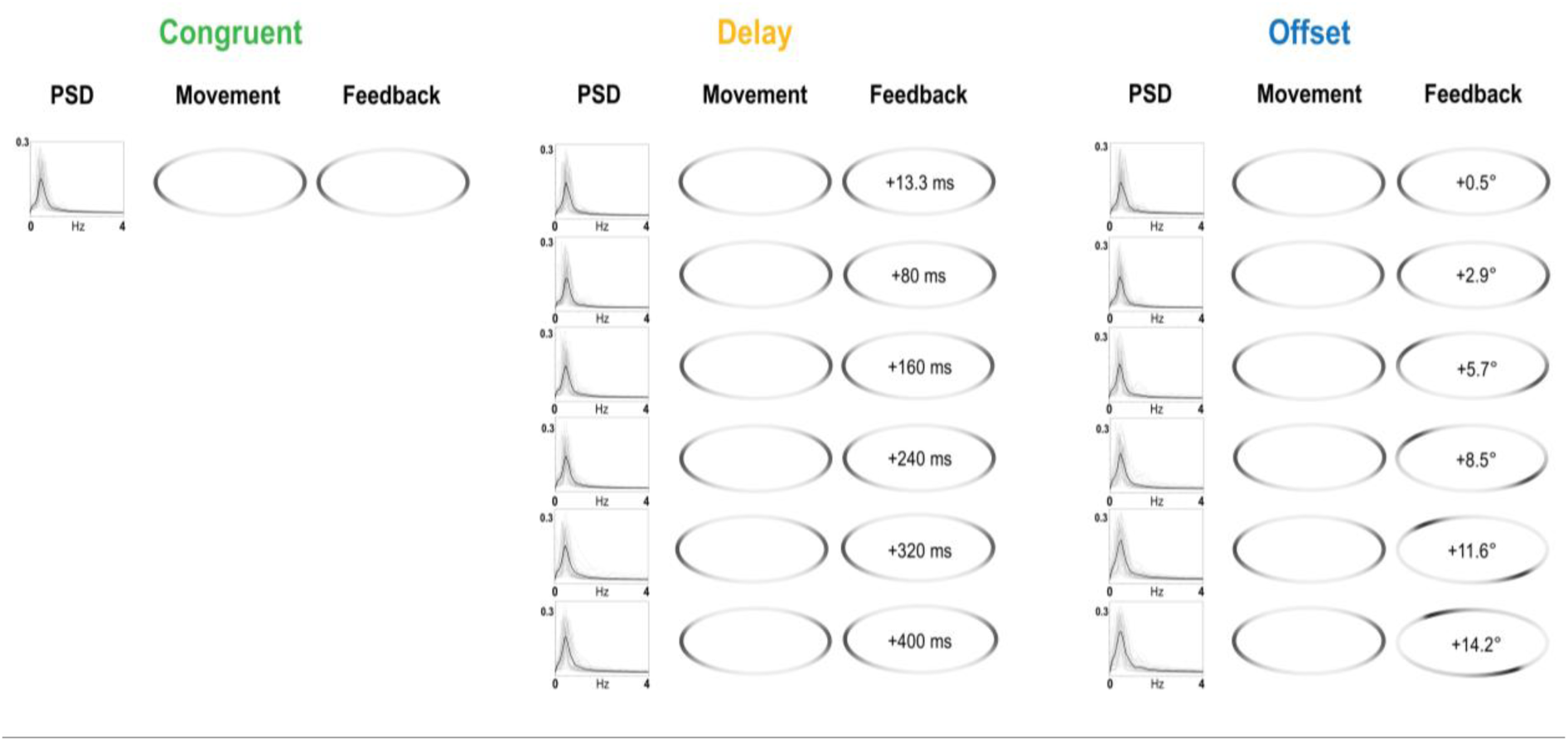
Kinematics of the executed movements and the visual movement feedback in each condition. The left plots in each condition show the averaged power spectral densities (PSD, cf. Figs. S2-S3). The elliptical plots show the averaged velocities of executed movements (left) and visual movement feedback (right) projected onto the elliptical trajectory (see Methods); darker parts indicate slower movement velocities. In all conditions, participants adhered to kinematic invariants described by the two-thirds power law; i.e., slowing down at stronger curvatures (left and right sides of the ellipse). The visual feedback also reflected this in the congruent (left) and the delayed (middle) conditions, the latter of which were effectively a simple lagging playback of the executed movements. In contrast, note the systematic shift of velocity profiles introduced by constant spatial offsets along the elliptical trajectory (right).

There were six different levels of spatial offset (0.5, 2.9, 5.7, 8.5, 11.6, and 14.2° visual angle). Crucially, for comparability of the six delay and offset levels, each level was matched in terms of the average Euclidean distance between the physical (pen) and displayed feedback (dot) positions on the Ellipse. I.e., for each delay level, we calculated the average Euclidean distance based on a geometrical model assuming constant velocity movement along the ellipse at 0.33 Hz. This was set as the displacement of the dot relative to the pen position in the respective “offset” levels.

The participants’ task was to decide, as quickly and correctly as possible, whether the visual feedback (dot movement) was congruent, delayed, or offset with respect to their own movement. The participants were instructed to move until they reached a decision, and then end each trial by pressing the Enter key. If no decision was reached within 20 s, the trial ended automatically. The participants then indicated whether the visual feedback was congruent (in German: “identisch”), delayed (“zeitlich verzögert”) or offset (“räumlich versetzt”), using the numeric keys 1-3 (assignment to the response options randomized across participants), and confirming again with the Enter key.

After about one week (7.12 ± 1.36 days, to avoid memory effects, cf. Knoblich & Prinz, 2001), participants completed the second, observation phase. This phase of the experiment served as a control condition for visual sensitivity in the absence of motor signals. The experimental setup with the covered tablet was identical to the first phase of the experiment; brown noise was also played throughout the entire phase. Participants viewed a playback of the recorded visual movement feedback of their movements performed in the first (execution) part in randomized order; no movement was executed (Fig. 1C). All playback trials were restricted to a maximum duration of 8 s, to reduce variability in trial length and, thus, to prevent participants inferring feedback conditions from decision time (i.e., trial length) alone. E.g., participants could have inferred a strong mismatch trial from a difference in trial length alone. It should be noted that, in these cases (about 60% of trials, see Supplement), participants had comparably shorter visual movement presentations in the observation than in the execution part. We therefore calculated a control analysis with matched trial lengths in both parts; i.e., limiting the analysis to only trials with a duration of <8s and excluding the other trials from the execution part as well. Note that this analysis can be seen as excluding trials with very high uncertainty; i.e., where participants took very long to reach a decision. We report all replications of the relevant analysis in the Supplementary material (Tables S7-S9, Fig. S5). As the resulting effects were nearly identical to those of our main analysis, in the main text, we only report where they differed from key significant results. After the presentation of each trial, participants indicated whether they believed the observed movement had been from a congruent, delayed, or offset movement trial (in a format identical to the execution phase).

In each phase of the experiment, participants completed 144 trials in total, with 144/3=48 trials per condition (congruent, delay, offset; all trials of the execution phase were included in the observation phase, albeit some were shortened, see above). Each of the 6 delay and offset levels was presented 8 times, interspersed with congruent trials, in randomized order. The 144 trials in each phase were divided into six blocks of 24 trials each. After completion of the experiment, the participants received compensation (10 € per hour or course credit).

### Movement data analysis

We analyzed the participants’ velocity and spectral profiles to evaluate the comparability of the executed movements. We first resampled the data at 75Hz, then epoched the trials according to the 13 experimental conditions. Trials in which participants made erroneous movements (i.e., clockwise, stopped for longer than 500 ms, or changed direction) were excluded from further data analysis; we also excluded trials in which the pen’s trajectory deviated more than 10% from the ideal elliptical trajectory. A stringent exclusion of these trials was necessary, because these deviations could immediately reveal clues about which condition the participant was in. With these criteria, we rejected 17.7% of all trials (Table S2); the number of excluded trials did not significantly differ across different feedback conditions (*χ*^2^(2)=1.58, *p*=.45, *Kendall’s W*=0.025) or the different stimulus levels (*χ*^2^ (12)=6.07, *p*=.912, *Kendall’s W*=0.016). Figure S1 shows the traces of all executed movements across conditions.

The average movement velocity was determined by dividing the number of total cycles per trial by the total time per trial; and averaging these values across trials per condition. We conducted a one-way repeated-measures analysis of variance (rmANOVA) to determine whether the average velocity differed between feedback conditions; and a 2×6 rmANOVA with the factors condition (offset, delay) and mismatch level (i.e.; the six respective levels of offset and delay) to determine whether it differed depending on mismatch levels. Furthermore, we looked at the participants’ velocity profiles; i.e., variations in movement speed along the elliptical trajectory, computed by the derivative form:

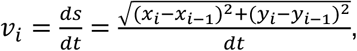

where *v_i_* is the estimated instantaneous speed of the i-th time sample. (***x_i_, y_i_***) represents the pen/cursor position at the i-th time point, and *dt* is the temporal distance between the two samples (i.e., for 75Hz = 0.013s). These estimates were smoothed using a Savitzky–Golay filter (15 time points window length, 0.2s) to reduce noise.

To visualize the velocity profiles (Fig. 2), we remapped the smoothed instantaneous velocity profiles from each valid drawing round onto our elliptical geometry template using a nearest-neighbor projection (see Supplementary material). The remapped velocity profiles of executed (pen) movements and visual feedback (dot movements) were averaged across completed elliptical cycles in each trial, then across trials per condition.

We also calculated the power spectral densities (PSD) of the pen movements in a range from 0-4Hz (cf. Foulkes & Miall, 2000), for movements along the x-axis and y-axis separately (cf. Supplementary material). The PSDs were tested for significant differences between conditions and mismatch levels using rmANOVAs as for the velocities.

### Analysis of detection and discrimination performance

To analyse the correct responses in the experimental phases, and their differences between conditions, we used GLMMs with binomial distribution and logit link function. I.e., we conducted a logistic regression with nominal-scale predictors, which we embedded in a hierarchical model, due to the two experimental parts. Level 1 represented the repeated measures within the subjects, while Level 2 represented the subject level. The Intraclass Correlation Coefficient (ICC) was 0.058. This indicates that only about 6% of the variance was due to differences between the test subjects; 94% of the variance was due to differences within individuals (e.g. experimental part, stimulus conditions and stimulus level; cf. Eid et al., 2017). The models could furthermore include other predictors like stimulus condition or stimulus level and their interactions. The nominal-scale predictors (experimental phase, condition, stimulus level) were dummy coded, with “execution phase” and “congruent” as the reference categories (“0”).

We quantified classification performance as the model-based mean probability of correctly identifying the type of visual movement feedback (congruent, delayed, or offset). Following our preregistered hypotheses, we calculated pairwise contrasts between the respective classification performances in the execution > observation phase for each stimulus condition. Additionally, we examined whether the influence of feedback condition depended on the experimental phase (i.e., interaction effect). To analyze the influence of the different mismatch levels (i.e., the 6 different levels of delay and offset, respectively) in the two parts, we calculated a second model with an explicit interaction between the predictors of the experimental phase and the stimulus level. The output of these analyses is a log odds ratio that is symmetric around zero (which corresponds to maximum uncertainty). I.e., a negative log odds ratio meant that a correct response in the execution phase was more likely than in the observation phase, whereas a positive log odds ratio would mean that a correct response was more likely during observation.

To determine whether classification performance was above chance level, we tested the model-based means against 1/3. Here, a positive or negative log odds ratio meant above- or below-chance level classification, respectively. Note that, in principle, participants could reach above chance level classification for delays (or offsets) while classifying similarly many, or even more delayed trials as offsets, and vice versa. Our three-alternative-choice response format allowed us to test, additionally, for *discriminative* performance between the two different kinds of visuomotor incongruence; i.e., whether participants identified the correct incongruence type more frequently than the incorrect one. We quantified this discriminative performance by calculating the ratio of the “temporal delay” or “spatial offset” responses to all incongruent responses, per stimulus level. We used a linear mixed model (LMM) with the proportion of correct responses as the dependent variable; and experimental phase (execution and observation) and stimulus conditions (delay and offset) as predictors. To analyze differences in discrimination performance at different mismatch levels, we calculated an additional LMM with the proportion of correct responses as the dependent variable and experimental phase (execution and observation) and mismatch level as predictors. We then determined model-based estimated marginal means, and calculated pairwise contrasts for the discrimination performances; and whether the observed proportions were significantly above chance level of 0.5. This tested our preregistered hypotheses H2 and H3 (see Table S1).

### Linear regressions

To test whether detection performance on the execution phase improved with increasing mismatch (delay or offset) level, we conducted linear regression analyses. To enable better comparability of the resulting regression weights (slopes), we normalised the predictor “stimulus level”; i.e., the 6 levels of delay and offset were normalized between 0 and 1, respectively. To test hypotheses H3, H4b, and H4c, we calculated a two-way rmANOVA with the factors experimental phase (execution, observation) and feedback condition (delay, offset) on the resulting regression weights for each participant. To test hypothesis H4a, we calculated an analogous two-way rmANOVA on the average detection performances. These analyses were calculated on the % trials in which participants gave the correct response (“classification”) and on the % trials in which participants correctly distinguished between delays and offsets when perceiving a visuomotor mismatch (“incongruence discrimination”).

### Reaction time analysis

To evaluate differences in reaction times across conditions, we calculated a one-way rmANOVA with the factor condition (congruent, offset, delay). As the reaction time distribution was not normal, a log transformation was applied across all participants. We calculated linear regressions for the delay and offset conditions separately (using normalized mismatch levels, as above), and used a paired-samples t-test to test for differences in slopes.

In all of the above ANOVAs, we tested for violations of the assumption of sphericity with Mauchly’s test, and applied Greenhouse-Geisser correction if required. In all of the above analyses, all post-hoc tests and pairwise comparisons (also against chance level) were Holm-Bonferroni-adjusted for multiple comparisons.

## Results

### Movement execution

Participants executed overall comparable movements across conditions, with the expected covariation of velocity and curvature (Fig. 2, Fig. S1). The average movement speed (0.4 Hz) did not differ significantly between conditions (*F*(2,10)=0.90, *p*=.44, η ^2^=0.15), but decreased somewhat with increasing mismatch levels (*F*(3.10,96.19)=8.35, *p*<.001, η ^2^=0.21, Table S3). At the highest mismatch levels (400ms, 14.2°), participants moved faster in the delay than in the offset condition (*t*(31)=4.42, *p*<.001, *d*=0.78). The PSDs showed clear peaks without higher-frequency components, suggesting regular movements (with overall lower power in the congruent condition, Figs. S2-S3).

### Detection and discrimination of delayed and offset visual movement feedback in the execution phase

During the execution phase, participants correctly (i.e., significantly above chance i.e. 33%) classified congruent feedback, delays >= 160 ms, and offsets >= 5.7° (Fig. 3A and Table 1, cf. Table S4). The lowest two delay and offset levels, respectively, were most frequently (incorrectly) classified as “congruent”. To test whether detection performance improved consistently with increasing mismatch (delay or offset) level, we conducted linear regression analyses. Indeed, there were significant differences in performance depending on the level of delay (*F*(1,190)=122.3, *p*<.001, *R^2^*=.40) and offset (*F*(1,190)=67.81, *p*<.001, *R^2^*=.26), respectively. The performance improvement with increasing amount of visual feedback incongruence could be linearly approximated for delays (β=0.66, *SE*=0.06, *t*=11.06, *p*<.001) and offsets (β=0.58, *SE*=0.07, *t*=8.24, *p*<.001, see Fig. 3B). We also tested for discriminative performance; i.e., whether participants could adequately distinguish delays from offsets in trials where they perceived visuomotor incongruence. Consistent with classification (see above), the average discriminative performance was significantly above chance (50%) for delays >= 160 ms, and offsets of 8.5 and 11.6° (with a statistical trend at 5.7 and 14.2°, Table 1). Discrimination did not change significantly with mismatch level (linear regressions revealed positive, but not significant slopes). On average, discriminative performance was significantly better for delays than for offsets (but the post-hoc contrasts at the individual mismatch levels did not reach significance, see Table S5).

**Figure 3.**
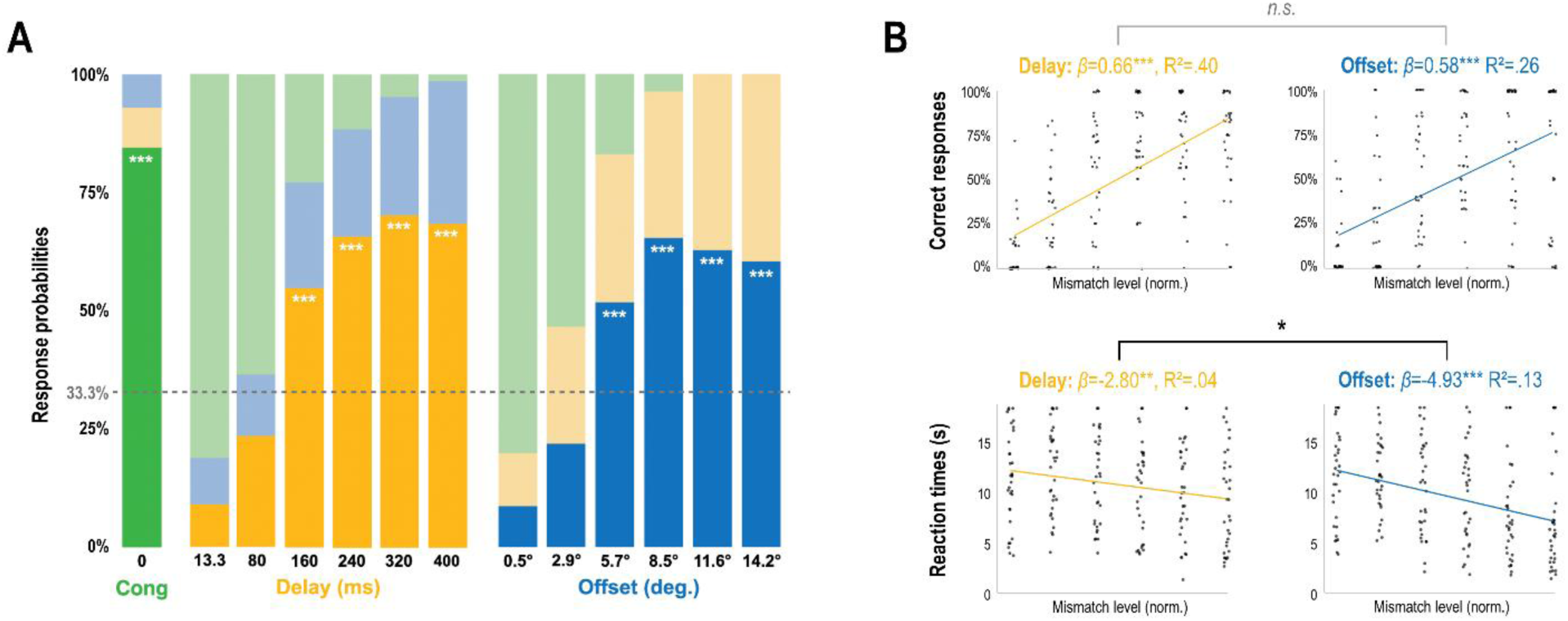
Response behavior in the execution phase. **A:** Each column shows the stacked model-based mean response probabilities in each condition (green=congruent, yellow=delayed, blue=offset). The correct response option is always displayed at the bottom and in a stronger saturation. The grey dashed line indicates the chance i.e. guessing level (33%). White asterisks denote correct responses significantly above chance level; \**p*<.05. \*\**p*<.01. \*\*\**p*<.001. **B:** Results of linear regressions between response behavior vs mismatch level in the delay and offset conditions, respectively. The mismatch (delay and offset) levels were normalized for better comparability of the regression weights, see Methods. The dots represent the individual participants’ averages. The top plots show non-zero positive linear relationships for correct responses (cf. panel A), which did not differ significantly between conditions. The bottom plots show non-zero negative linear relationships for reaction times, which were significantly stronger for offsets than for delays. See Table 1 for details.

**Table 1.** Mean rating performances (model-based means) and reaction times in the execution phase, with associated standard errors of the mean in brackets. Classification quantifies the probability of correct (congruent/delay/offset) responses; incongruence discrimination quantifies the proportion of correctly distinguished delays vs offsets in incongruent trials.

|  | Classification |  | Incongruence discrimination |  | Reaction time (s) |
| --- | --- | --- | --- | --- | --- |
| | Prob. correct | $p >$<br>chance (33.3%) | Prob. correct | $p >$<br>chance (50%) | |
| Congruent |  |  |  |  |  |
|  | 0.86 (0.01) | <.0001 | - | - | 11.67 (0.75) |
| Delay |  |  |  |  |  |
| 13.3 ms | 0.08 (0.02) | >.99 | 0.45 (0.08) | .74 | 11.66 (0.80) |
| 80 ms | 0.23 (0.03) | >.99 | 0.63 (0.07) | .05 | 12.48 (0.75) |
| 160 ms | 0.56 (0.04) | <.0001 | 0.73 (0.07) | <.001 | 11.41 (0.77) |
| 240 ms | 0.68 (0.04) | <.0001 | 0.75 (0.06) | <.001 | 10.6 (0.81) |
| 320 ms | 0.72 (0.04) | <.0001 | 0.75 (0.06) | <.001 | 10.2 (0.81) |
| 400 ms | 0.71 (0.04) | <.0001 | 0.73 (0.06) | <.001 | 9.24 (0.86) |
| Mean | 0.5 (0.03) | <.0001 | 0.69 (0.04) | <.0001 | 10.96 (0.72) |
| Offset |  |  |  |  |  |
| 0.5° | 0.08 (0.02) | >.99 | 0.32 (0.08) | >.99 | 11.66 (0.80) |
| 2.9° | 0.21 (0.03) | >.99 | 0.41 (0.07) | >.99 | 11.72 (0.62) |
| 5.7° | 0.53 (0.04) | <.0001 | 0.64 (0.06) | .05 | 10.83 (0.84) |
| 8.5° | 0.66 (0.04) | <.0001 | 0.69 (0.06) | .01 | 9.83 (0.83) |
| 11.6° | 0.64 (0.04) | <.0001 | 0.65 (0.06) | <.05 | 7.95 (0.84) |
| 14.2° | 0.62 (0.04) | <.0001 | 0.64 (0.06) | .05 | 7.17 (0.88) |
| Mean | 0.46 (0.03) | <.0001 | 0.58 (0.04) | .02 | 9.84 (0.69) |

Significant p-values are in bold font. All p-values are Holm-Bonferroni adjusted for multiple comparisons.

### Reaction times

The participants’ reaction times (Table 1) differed significantly between feedback conditions (rmANOVA, *F*(1.56,48.29)=9.92, *p*<.001, η ^2^=0.24). Participants responded, on average, faster in the offset condition than in the delayed (*t*(31)=-3.535, *p*<.01, *d*=0.63) or congruent (post-hoc *t*(31)=3.61, *p*<.01, *d*=0.64) conditions. Linear regressions (Fig. 3C) revealed that participants required consistently less time to reach a decision with increasing delay (*F*(1,190)=8.796, *p*<.01, *R^2^*=.04; β=-2.8, *SE*=0.94, *t*=-2.97, *p*<.01) and offset (*F*(1,190)=27.62, *p*<.001, *R^2^*=0.13; β=-4.93, *SE*=0.93, *t*=-5.26, *p*<.001). A paired t-test on the mean regression slopes (not preregistered) furthermore showed that this tendency was significantly stronger for offset than for delayed visual movement feedback (*t*(31)=3.62, *p*<.01, *d*=0.64). In other words, while reaction times during very low delay and offset trials were comparable, they decreased more strongly when the amount of offset increased.

### Detection of delayed vs offset visual movement feedback in the observation phase

Classification performance in the observation phase was overall closer to chance level than during execution (Fig. 4A; cf. Fig. S4). Note that due to the principal indistinguishability of delayed and congruent playback trials (see Methods), the comparison with the delay condition during execution is trivial. I.e., unsurprisingly, classification performance in the delay condition was not different from chance level; overall responses were practically identical with those in the congruent condition (Table 2).

**Figure 4.**
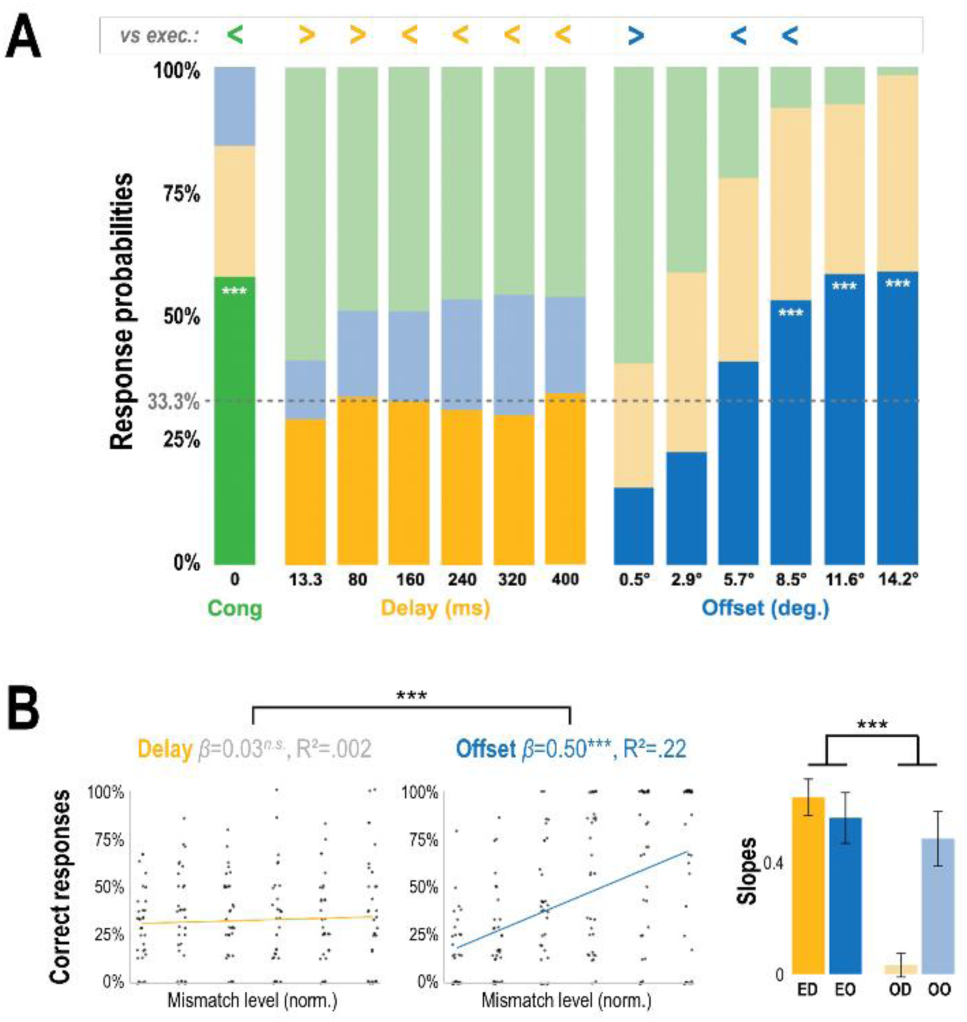
**A:** Response probabilities in the observation phase. Format as in Figure 3A. The inlay above the bars indicates significant differences against execution; i.e. “<” means poorer performance during observation than during execution; “>” means the converse. See Table 2 for details. **B:** Results of linear regressions between correct detection performance vs mismatch level in the delay and offset conditions, respectively. The bar plots show the mean regression weights with associated standard errors. An rmANOVA revealed significantly steeper slopes during execution (main effect of phase; but the post-hoc contrast only reached significance for delays, not offsets).

**Table 2.**
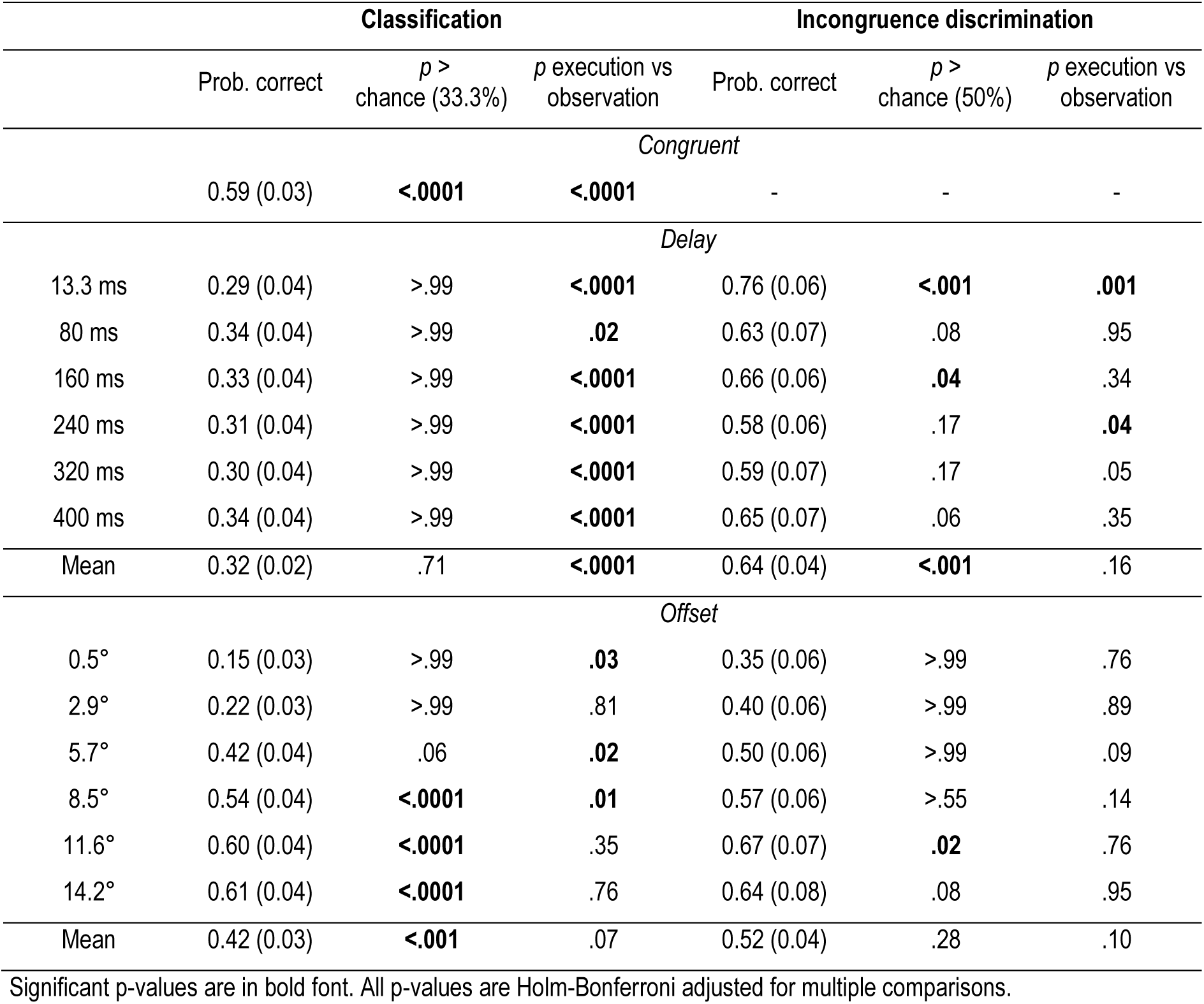
Mean rating performances in the observation phase, with associated standard errors of the mean in brackets, and comparisons against the execution phase (cf. Table 1).

|  | Classification |  |  | Incongruence discrimination |  |  |
| --- | --- | --- | --- | --- | --- | --- |
| | Prob. correct | $p >$<br>chance (33.3%) | $p$ execution vs<br>observation | Prob. correct | $p >$<br>chance (50%) | $p$ execution vs<br>observation |
| *Congruent* |  |  |  |  |  |  |
|  | 0.59 (0.03) | **<.0001** | **<.0001** | - | - | - |
| *Delay* |  |  |  |  |  |  |
| 13.3 ms | 0.29 (0.04) | >.99 | **<.0001** | 0.76 (0.06) | **<.001** | **.001** |
| 80 ms | 0.34 (0.04) | >.99 | **.02** | 0.63 (0.07) | .08 | .95 |
| 160 ms | 0.33 (0.04) | >.99 | **<.0001** | 0.66 (0.06) | **.04** | .34 |
| 240 ms | 0.31 (0.04) | >.99 | **<.0001** | 0.58 (0.06) | .17 | **.04** |
| 320 ms | 0.30 (0.04) | >.99 | **<.0001** | 0.59 (0.07) | .17 | .05 |
| 400 ms | 0.34 (0.04) | >.99 | **<.0001** | 0.65 (0.07) | .06 | .35 |
| Mean | 0.32 (0.02) | .71 | **<.0001** | 0.64 (0.04) | **<.001** | .16 |
| *Offset* |  |  |  |  |  |  |
| 0.5° | 0.15 (0.03) | >.99 | **.03** | 0.35 (0.06) | >.99 | .76 |
| 2.9° | 0.22 (0.03) | >.99 | .81 | 0.40 (0.06) | >.99 | .89 |
| 5.7° | 0.42 (0.04) | .06 | **.02** | 0.50 (0.06) | >.99 | .09 |
| 8.5° | 0.54 (0.04) | **<.0001** | **.01** | 0.57 (0.06) | >.55 | .14 |
| 11.6° | 0.60 (0.04) | **<.0001** | .35 | 0.67 (0.07) | **.02** | .76 |
| 14.2° | 0.61 (0.04) | **<.0001** | .76 | 0.64 (0.08) | .08 | .95 |
| Mean | 0.42 (0.03) | **<.001** | .07 | 0.52 (0.04) | .28 | .10 |

Significant p-values are in bold font. All p-values are Holm-Bonferroni adjusted for multiple comparisons.

For the offset condition, there was a non-significant tendency for better classification during execution than observation (log-odds difference=-0.15, *SE*=0.08, *z*=1.82, *p*=.07, Table 2, cf. Table S6). Notably, participants showed significantly better classification during execution than observation at medium offsets of 5.7 and 8.5° (Table 2). This difference, although still present, did not reach significance when matching trial lengths in both parts (control analysis, see Table S8). At higher offsets (11.6, 14.2°), performance was also better during execution than observation, but the differences did not reach significance. At the smallest offset level (0.5°), classification performance was significantly poorer during execution than observation; however, this difference became non-significant after matching trial lengths in both parts (Table S8). Furthermore, there was a tendency for an overall better discrimination in the offset condition during execution than observation, but none of the post-hoc contrasts reached significance (Table 2).

Linear regressions on the detection slopes (see above) revealed a significant positive relationship for offsets but not for delays, see Figure 4B. A rmANOVA with the factors experimental phase (execution, observation) and feedback condition (delay, offset) on the resulting regression weights revealed a significant main effect (*F*(1,31)=42.38, *p*<.001 η ^2^ = 0.58), with steeper slopes in the execution than in the observation phase; and a significant interaction (*F*(1,31)=21.83, *p*<.001, η ^2^ = 0.41). Post-hoc t-tests revealed significantly steeper slopes during the execution than the observation part for delays (*t*(31)=7.995, *p*<.001, *d* = 1.41) but not for offsets (*t*(31)=0.946, *p*=.78, *d* = 0.17). See Figure 4B. An analogous rmANOVA on discrimination performance yielded no significant difference between execution and observation (main effect, *F*(1, 31) = 0.38, *p* = .54, η ^2^ = 0.01).

## Discussion

Here, we tested for commonalities and differences in the detection of visuomotor incongruence arising from delays vs offsets, using an elliptical task that allowed to match movement trajectories and to control for the presence of spatial discrepancy. Our key findings were (1) a similar perceptual sensitivity to delays and offsets during execution; and (2) a significantly and selectively improved detection of certain spatial offsets during execution compared with observation.

Firstly, the participants’ responses suggest overall similar perceptual sensitivities to visuomotor mismatches resulting from delaying or offsetting visual feedback, when those manipulations are matched in terms of their trajectory and average spatial discrepancy. Matched levels of delay and offset were comparably frequently correctly classified, and comparably frequently correctly distinguished from each other. Specifically, participants were able to correctly identify mismatches greater than 160 ms or 5.7° as delays or offsets, respectively. Furthermore, detection performance increased significantly with increasing delay and offset, while reaction times decreased, supporting our first key hypothesis. Very low mismatch levels were mostly classified as congruent, suggesting they fell below perceptual thresholds very similar to those identified in previous work (e.g. Leube et al., 2003; Farrer et al., 2008; Krugwasser et al., 2019).

These results suggest that mismatching visual movement feedback arising from delays (with asynchronous kinematics) and offsets (with synchronous kinematics but violating biological motion laws) may be perceptually equally salient. Thus, they tentatively support previous suggestions of comparable perceptual thresholds for spatial and temporal action feedback manipulations (e.g., Farrer et al., 2008; Krugwasser et al., 2019).

Notably, however, the overall discriminative performance was significantly better for delays than for offsets (although the post-hoc tests of differences at individual mismatch levels did not reach significance). I.e., participants somewhat more frequently confused offsets with delays, than vice versa. Conversely, participants responded significantly faster when reporting offsets than delays. This could imply a speed-accuracy trade-off: While participants detected manipulated visual feedback more quickly when it violated biological laws (perhaps due to the sensitivity of the visual system to such violations, see Introduction), they could not tell as accurately in what way that feedback deviated from their movements. Interestingly, at very small offset vs delay levels the reaction time benefit was absent—and the discriminative performance was particularly poor for reported offsets compared with delays. In this light, an asynchrony of visual and motor kinematics (i.e., in our delay condition) could be seen as a more salient agency cue, based on a more precise visuomotor evaluation particularly at very small mismatch levels. While speculative, this interpretation speaks to previous proposals emphasizing the importance of temporal prediction for agency and self-other distinction (Rohde & Ernst, 2016; Haggard, 2017).

Our second key finding was that the detection of some (but not all) delays and offsets was improved during action compared to mere observation. Delayed playback trials were classified virtually at chance level. While discrimination was partly above chance level in the observe-delay condition, this can be interpreted as simply “not detecting offsets”. I.e., the overall response pattern was virtually identical with the congruent condition; showing that delayed playback was, in the absence of motor reference signals, indistinguishable from the playback of congruent trials. In contrast, offset playback trials were reliably detected during passive observation alone; suggesting that participants were *perceptually* sensitive to violations of biological kinematics (velocity-curvature covariation). This supports previous findings that the visual system is sensitive to nonbiological movements (Johansson, 1973; Viviani & Stucchi, 1992; de’Sperati & Stucchi, 1995; Dayan et al., 2007; Salomon et al., 2016; Fraser et al., 2025; Rolfs et al., 2025).

Crucially, action significantly enhanced the correct detection at medium-high offsets (5.7/8.5°). A similar tendency was observable for discriminative performance at these offset levels, which was significantly above chance level during execution, but not during observation (however, these differences did not reach significance). This partly supported our second key hypothesis. As performance plateaued at larger offsets, motor signals did not seem to provide any additional benefit once the visual velocity profile became too biologically implausible. It should be noted that this difference did not reach significance when analysing only trials with comparable lengths (which could, however, have resulted from the reduced trial numbers in the control analysis). Thus, these medium-high offset levels potentially constituted a critical threshold for visuomotor self-other distinction. Together with the (albeit non-significantly) steeper slopes of classification performance during execution compared with observation, this tentatively suggests action led to a ‘sharper’ differentiation between congruent and offset visual movement feedback.

The poorer detection performance for very low mismatch levels during execution compared with observation may seem counter-intuitive. However, it likely resulted from a tendency to guess more frequently during the observation of offsets that were not clearly perceptually distinguishable in the absence of (motor) reference signals. This tendency would have yielded higher frequencies of “correct” responses than during execution, where participants were incorrectly, but systematically rating the visual movement feedback at small mismatch levels as congruent. This interpretation is supported by our control analysis, which showed that the performance during observation of 0.5° offsets dropped to a level not significantly different from the respective execution condition when controlling for the different number of potential high-uncertainty trials (i.e., with unusually long reaction times). Alternatively, during execution, the movement task could have competed for cognitive-attentional resources with the detection task, which could have impaired the detection of small mismatches in particular. Disentangling between these alternative explanations is a key question for future work. Assessments of confidence could help clarify this; i.e., by revealing potential guessing tendencies in the observation part. Furthermore, the isolation of motor from ‘non-motor’ (e.g., somatosensory) movement signals to mismatch detection needs to be addressed by future work through inclusion of passive movement controls. Finally, there were small but significant influences of mismatches on the executed movements; while these were balanced across our execution and observation parts, future work should attempt better control.

In conclusion, our results suggest remarkably similar perceptual thresholds for visuomotor mismatches resulting from asynchronous vs nonbiological visual kinematics; and that the perception of these mismatches, even of some of those resulting from nonbiological kinematics, may be improved by action.

## Declarations

### Ethics approval

The experiment was approved by the ethics committee of the University Medicine of Greifswald.

### Consent to participate

All participants signed written informed consent.

### Consent for publication

All participants signed written informed consent.

### Availability of data and materials

The data will be made available upon request. The data are not available publicly due to ethical and privacy restrictions.

### Code availability

The code will be made available upon request.

## Acknowledgments

We thank Eric Kühn of the science workshop of the Universität Greifswald for 3D-printing the elliptical guide; Josephine Gräfe, Luka Henkel, Eric Bendt and Cathia Lindloff for their help with data collection; Linnea Hertel for creating the setup figure; and Matti Lyko and Michael Höhle (Statistical Consulting, University of Greifswald) for statistical consulting. We also thank our anonymous reviewers for their many helpful suggestions.

## References

Bidet-Ildei, C., Méary, D., & Orliaguet, J.-P. (2008). Visual preference for isochronic movement does not necessarily emerge from movement kinematics: A challenge for the motor simulation theory. Neuroscience Letters, 430(3), 236–240. 10.1016/j.neulet.2007.10.040

Dayan, E., Casile, A., Levit-Binnun, N., Giese, M. A., Hendler, T., & Flash, T. (2007). Neural representations of kinematic laws of motion: Evidence for action-perception coupling. Proceedings of the National Academy of Sciences, 104(51), 20582–20587. 10.1073/pnas.0710033104

de’Sperati, C., & Stucchi, N. (1995). Visual tuning to kinematics of biological motion: The role of eye movements. Experimental Brain Research, 105(2). 10.1007/BF00240961

Farrer, C., & Frith, C. D. (2002). Experiencing Oneself vs Another Person as Being the Cause of an Action: The Neural Correlates of the Experience of Agency. NeuroImage, 15(3), 596–603. 10.1006/nimg.2001.1009

Farrer, C., Bouchereau, M., Jeannerod, M., & Franck, N. (2008). Effect of distorted visual feedback on the sense of agency. Behavioural Neurology, 19(1–2), 53–57.

Foulkes, A. J. M., & Miall, R. C. (2000). Adaptation to visual feedback delays in a human manual tracking task. Experimental brain research, 131, 101–110.

Fraser, D. S., Cook, J., & Di Luca, M. (2025). Biological kinematics: A detailed review of the velocity-curvature power law calculation. Experimental Brain Research, 243(5), 107. 10.1007/s00221-025-07065-0

Frith, C. D., Blakemore, S.-J., & Wolpert, D. M. (2000). Abnormalities in the awareness and control of action. Philosophical Transactions of the Royal Society of London. Series B: Biological Sciences, 355(1404), 1771–1788. 10.1098/rstb.2000.0734

Green, P. & MacLeod, C.J. (2016). SIMR: an R package for power analysis of generalized linear mixed models by simulation. Methods Ecol Evol, 7, 493–498. 10.1111/2041-210X.12504

Haggard, P. (2017). Sense of agency in the human brain. Nature Reviews. Neuroscience, 18(4), 196–207. 10.1038/nrn.2017.14

Huh, D., & Sejnowski, T. J. (2015). Spectrum of power laws for curved hand movements. Proceedings of the National Academy of Sciences, 112(29). 10.1073/pnas.1510208112

Johansson, G. (1973). Visual perception of biological motion and a model for its analysis. Perception & Psychophysics, 14(2), 201–211. 10.3758/BF03212378

Jordan, M. I., & Rumelhart, D. E. (1992). Forward models: Supervised learning with a distal teacher. Cognitive Science, 16(3), 307–354. 10.1207/s15516709cog1603_1

Knoblich, G., & Prinz, W. (2001). Recognition of self-generated actions from kinematic displays of drawing. Journal of Experimental Psychology: Human Perception and Performance, 27(2), 456–465. 10.1037/0096-1523.27.2.456

Knoblich, G., Seigerschmidt, E., Flach, R., & Prinz, W. (2002). Authorship effects in the prediction of handwriting strokes: Evidence for action simulation during action perception. The Quarterly Journal of Experimental Psychology Section A, 55(3), 1027–1046. 10.1080/02724980143000631

Krugwasser, A. R., Harel, E. V., & Salomon, R. (2019). The boundaries of the self: The sense of agency across different sensorimotor aspects. Journal of Vision, 19(4), 14. 10.1167/19.4.14

Lacquaniti, F., Terzuolo, C., & Viviani, P. (1983). The law relating the kinematic and figural aspects of drawing movements. Acta Psychologica, 54(1–3), 115–130. 10.1016/0001-6918(83)90027-6

Leube, D. T., Knoblich, G., Erb, M., Grodd, W., Bartels, M., & Kircher, T. T. (2003). The neural correlates of perceiving one’s own movements. NeuroImage, 20(4), 2084–2090. 10.1016/j.neuroimage.2003.07.033

Limanowski, J., Kirilina, E., & Blankenburg, F. (2017). Neuronal correlates of continuous manual tracking under varying visual movement feedback in a virtual reality environment. NeuroImage, 146, 81–89. Limanowski, 10.1016/j.neuroimage.2016.11.009

Limanowski, J. (2022). Precision control for a flexible body representation. Neuroscience & Biobehavioral Reviews, 134, 104401.

Limanowski, J. (2026). Cortical candidates for self-other distinction based on visual and action cues: Where do we stand?. Brain Structure and Function, 231(1), 10.

Miall, R. C., Weir, D. J., Wolpert, D. M., & Stein, J. F. (1993). Is the Cerebellum a Smith Predictor? Journal of Motor Behavior, 25(3), 203–216. 10.1080/00222895.1993.9942050

Miall, R. C., & Wolpert, D. M. (1996). Forward Models for Physiological Motor Control. Neural Networks, 9(8), 1265–1279. 10.1016/S0893-6080(96)00035-4

Peirce, J., Gray, J. R., Simpson, S., MacAskill, M., Höchenberger, R., Sogo, H., Kastman, E., & Lindeløv, J. K. (2019). PsychoPy2: Experiments in behavior made easy. Behavior research methods, 51(1), 195–203. 10.3758/s13428-018-01193-y

Rohde, M., & Ernst, M. O. (2016). Time, agency, and sensory feedback delays during action. Current Opinion in Behavioral Sciences, 8, 193–199. 10.1016/j.cobeha.2016.02.029

Rolfs, M., Schweitzer, R., Castet, E., Watson, T. L., & Ohl, S. (2025). Lawful kinematics link eye movements to the limits of high-speed perception. Nature Communications, 16(1), 1–17.

Salomon, R., Lim, M., Kannape, O., Llobera, J., & Blanke, O. (2013). “Self pop-out”: agency enhances self-recognition in visual search. Experimental brain research, 228(2), 173–181.

Salomon, R., Goldstein, A., Vuillaume, L., Faivre, N., Hassin, R. R., & Blanke, O. (2016). Enhanced discriminability for nonbiological motion violating the two-thirds power law. Journal of Vision, 16(8), 12. 10.1167/16.8.12

Stock, A.-K., Wascher, E., & Beste, C. (2013). Differential Effects of Motor Efference Copies and Proprioceptive Information on Response Evaluation Processes. PLoS ONE, 8(4), e62335. 10.1371/journal.pone.0062335

Synofzik, M., Vosgerau, G., & Newen, A. (2008). Beyond the comparator model: a multifactorial two-step account of agency. Consciousness and cognition, 17(1), 219–239.

Tanaka, T., & Imamizu, H. (2025). Sense of agency for a new motor skill emerges via the formation of a structural internal model. Communications Psychology, 3(1), 70.

Torricelli, F., Tomassini, A., Pezzulo, G., Pozzo, T., Fadiga, L., & D’Ausilio, A. (2023). Motor invariants in action execution and perception. Physics of Life Reviews, 44, 13–47. 10.1016/j.plrev.2022.11.003

Tsakiris, M., Haggard, P., Franck, N., Mainy, N., & Sirigu, A. (2005). A specific role for efferent information in self-recognition. Cognition, 96(3), 215–231. 10.1016/j.cognition.2004.08.002

Tsakiris, M., & Haggard, P. (2005). Experimenting with the acting self. Cognitive Neuropsychology, 22(3–4), 387–407. 10.1080/02643290442000158

Viviani, P., & Stucchi, N. (1992). Biological movements look uniform: Evidence of motor-perceptual interactions. Journal of Experimental Psychology: Human Perception and Performance, 18(3), 603–623. 10.1037/0096-1523.18.3.603

Wen, W., Brann, E., Di Costa, S., & Haggard, P. (2018). Enhanced perceptual processing of self-generated motion: Evidence from steady-state visual evoked potentials. NeuroImage, 175, 438–448. 10.1016/j.neuroimage.2018.04.019

Wen, W., & Imamizu, H. (2022). The sense of agency in perception, behaviour and human– machine interactions. Nature Reviews Psychology, 1(4), 211–222. 10.1038/s44159-022-00030-6

Wolpert, D.M., Miall, R.C., 1996. Forward models for physiological motor control. Neural Networks, 9, 1265–1279.

