## Supplementary Material for "Detection of mismatching visual kinematics during visuomotor incongruence: A comparison of spatially matched delays vs offsets along identical trajectories"

---

**Table S1. Preregistered hypotheses**

This table lists our preregistered hypotheses (https://osf.io/39tch). Please note that in the manuscript, following recommendations from our reviewers, we have condensed these hypotheses; i.e., we have omitted hypotheses centering on the (trivial) comparison of delay detection between execution and observation phases.

| Nr. | Hypothesis | Supported? | Relevant Result |
|---|---|---|---|
| H1 | The detection of incongruent visual movement feedback is better in the active than in the passive part (main effect). | Partially | Significant differences between classification during execution vs observation for congruent and delayed trials; and for certain levels of offsets. (Figure 4A, Table 2) |
| H2 | The discrimination among the three types of visual movement feedback is better in the active than in the passive part (main effect). | Partially | Significantly better detection of congruent and delayed movements during execution than observation. Above-chance level discrimination of delays vs offsets during execution but not observation. (Figure 4A, Table 2) |
| H2a | When moving, participants can distinguish between congruent, delayed, and offset visual action feedback above chance-level (active part). | Yes | Significant main effect classification and discrimination for each feedback type during execution. (Table 1) |
| H3 | When moving, participants show better detection and discrimination performance with increasing amount of visual action feedback incongruence (active part). | Partially | Significantly positive slopes in the regression analysis of delays and offsets. (Figure 3B)<br>Positive (but not significantly different from zero) slopes for discrimination between delays vs offsets. |
| H4a | In the passive part, participants are relatively better at detecting spatial offset than temporal delay, due to violations of the two-thirds power law in the replayed offset movements (interaction effect, cf. H1). | Yes | Chance-level detection of delays during observation. (Figure 4A, Table 2) |
| H4b | The hypothesized positive association between the amount of incongruence and detection performance (cf. H3) is stronger in the active than in the passive part (main effect). | Partially | More positive regression weights (slopes) for delays and offsets during execution compared with observation, but post-hoc contrast only significant for delays. (Figure 4B) |
| H4c | The hypothesized positive association between the amount of incongruence and detection performance (cf. H3) is stronger for offset than delay in the passive than in the active part (interaction effect). | Yes | Significant interaction effect on the regression weights. (Figure 4B) |

**Box S1: The Two-Thirds Power Law.**

Movements along curved paths are characterised by a specific „kinematic invariant" (Torricelli et al., 2022): The two-thirds power law.

This two-thirds power law is based on Binet and Courtier's 1893 observation that curves with a smaller radius of curvature are drawn more slowly than those with a larger radius (cited in Fraser et al., 2024). Formalised in 1983, this law describes a power function relating instantaneous tangential velocity to the radius of curvature of movement (Lacquaniti et al., 1983):

$$V(t) = kR(t)^{1-\beta}$$

V(t) is the tangential velocity, R(t) is the radius of curvature as a function of time, k is a velocity amplification factor, and β is an exponent with a value of approximately 1/3 (Bidet-Ildei et al., 2008). This means that the velocity of a movement increases along a less curved path and decreases along more curved sections (Salomon et al., 2016). This law is considered universal for biological movements and is firmly established in movement research and kinematics (Fraser et al., 2024). However, the exponent β is not a universal constant; rather, it results from the angular frequency of the trajectory shapes in question: For elliptical trajectories (angular frequency = 2), the model predicts β = 0.33, which corresponds to the two-thirds power law (Hun & Sejnowski, 2015). Shapes with lower angular frequencies (such as spirals) have higher exponents, while shapes with higher angular frequencies (such as rounded polygons) have lower exponents.

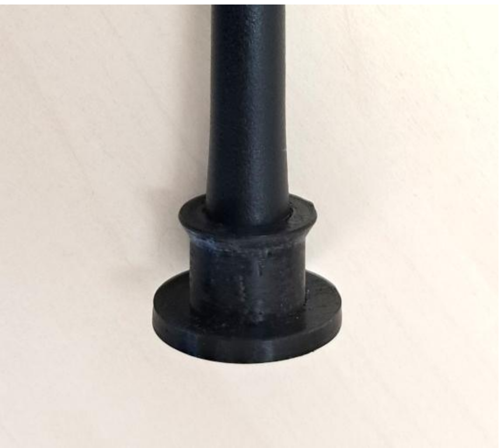

**Detail of pen in holder.**

The bottom part of the 3D printed, inverted-T shaped holder moved under the 3D printed ellipse. The pen was pressed firmly into the holder, and could only be removed with great force. This prevented the pen from being lifted during movement.

**Supplementary Methods: Determining spatial offsets along the elliptical trajectory**

The target ellipse was discretized into 500 points ***($x_i$, $y_i$)***, i=1,…,500. For each of these pen positions, we aimed to determine a corresponding cursor position on the same ellipse, located at a fixed Euclidean distance ***d*** behind the pen along the trajectory. This distance ***d*** represents the predefined spatial lag. However, the pre-sampled 500 coordinates on the ellipse, although sufficient for capturing the spatial resolution of the pen on the pad, were not fine enough to reliably identify the cursor positions at the exact spatial lag ***d***. For instance, with ***d*** = 0.006 and a given pen position ***($x_i$, $y_i$)***, the lagged cursor point might happen to fall outside of the 500 pre-sampled coordinates on the ellipse. Therefore, we implemented a numerical approach to estimate cursor positions that continuously spatially lag behind the subject-controlled pen positions along a predefined elliptical trajectory. To address this, we resampled the target ellipse at a much higher resolution with 20,000 points to generate a high dense trajectory ***($x^h_j$,$y^h_j$)***, j=1,…,20000. For each pen position on the target ellipse ***($x_i$, $y_i$)***, we then searched on this high-resolution ellipse for two candidate points whose Euclidean distance from the pen position matched the predefined spatial lag ***d*** (with tolerance < 1e-5).

To identify the correct lagging candidate, we used a motion vector-based directionality test:

1. Based on the predefined 500 points on the target ellipse, we estimated a motion vector for each pen position ***($x_i$, $y_i$)*** by using the previous point ***($x_{i-1}$, $y_{i-1}$),*** assuming counterclockwise movement: $Vmotion(i) = (x_i - x_{i-1}, y_i - y_{i-1})$ .
2. We computed vectors from the current pen position ***($x_i$, $y_i$)*** to each of the two candidate cursor positions on the ellipse.
3. We then calculated the dot product between each of these vectors and the motion vector $Vmotion(i)$
4. Finally, the candidate with a negative dot product was identified as the “offset” point, i.e. opposite (clockwise) to the direction of movement along the elliptical trajectory.

In sum, for each of the 500 pre-defined pen positions on the target ellipse, we identified the corresponding cursor position, which is located at a fixed spatial lag Euclidean distance ***d***, by searching a high-resolution resampling of the same ellipse (20,000 points). Among the candidate points at the correct distance, the lagging point was selected based on movement direction, determined via a dot product comparison with the local motion vector. This resulted in 500 paired positions between the pen and the spatially lagged cursor along the elliptical trajectory. As an example of numerical accuracy, for the smallest given spatial lag (***d*** = 0.006), the maximum relative error between the computed and proposed Euclidean distance was approximately 0.15%, which is visually negligible. For the largest lag distance (***d*** = 0.184), the relative error was approximately 0.003%.

**Table S2: Mean percentage of valid trials per condition**

| | Mean % | SD |
|---|---|---|
| | *Congruent* | |
| 0 | 82 | 38 |
| | *Delay* | |
| 13.3 ms | 83 | 37 |
| 80 ms | 82 | 39 |
| 160 ms | 84 | 36 |
| 240 ms | 81 | 39 |
| 320 ms | 83 | 38 |
| 400 ms | 83 | 37 |
| Mean | 82 | 39 |
| | *Offset* | |
| 0.5° | 80 | 40 |
| 2.9° | 82 | 38 |
| 5.7° | 84 | 37 |
| 8.5° | 81 | 39 |
| 11.6° | 80 | 40 |
| 14.2° | 82 | 38 |
| Mean | 83 | 38 |

**Figure S1: Executed pen movements in each condition**

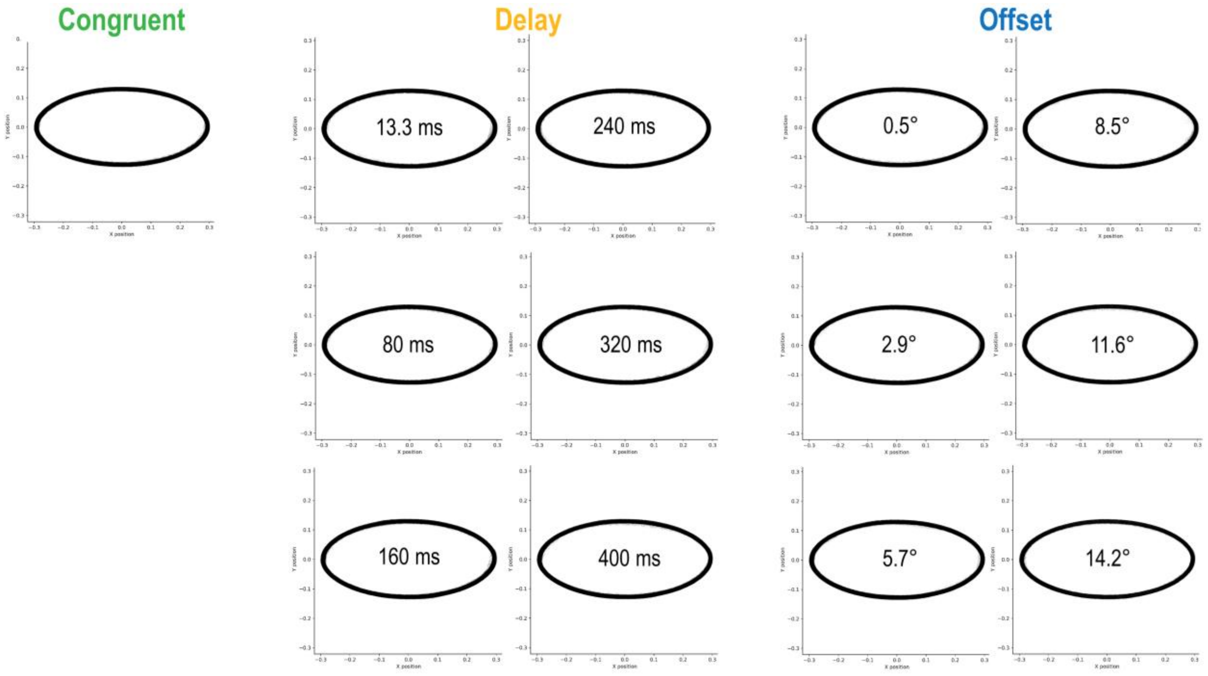

**Table S3: Average movement velocities (Hz)**

| | *Frequency* | *SD* | *SE* |
|---|---|---|---|
| | *Congruent* | | |
| 0 | 0.42 | 0.1 | 0.02 |
| | *Delay* | | |
| 13.3 ms | 0.41 | 0.09 | 0.02 |
| 80 ms | 0.42 | 0.1 | 0.02 |
| 160 ms | 0.41 | 0.1 | 0.02 |
| 240 ms | 0.4 | 0.1 | 0.02 |
| 320 ms | 0.38 | 0.11 | 0.02 |
| 400 ms | 0.38 | 0.09 | 0.02 |
| Mean | 0.4 | 0.1 | 0.01 |
| | *Offset* | | |
| 0.5° | 0.42 | 0.1 | 0.02 |
| 2.9° | 0.42 | 0.1 | 0.02 |
| 5.7° | 0.4 | 0.11 | 0.02 |
| 8.5° | 0.39 | 0.1 | 0.02 |
| 11.6° | 0.39 | 0.12 | 0.02 |
| 14.2° | 0.37 | 0.11 | 0.02 |
| Mean | 0.4 | 0.11 | 0.01 |

**Figure S2: Power spectral densities for the x- and y-axes**

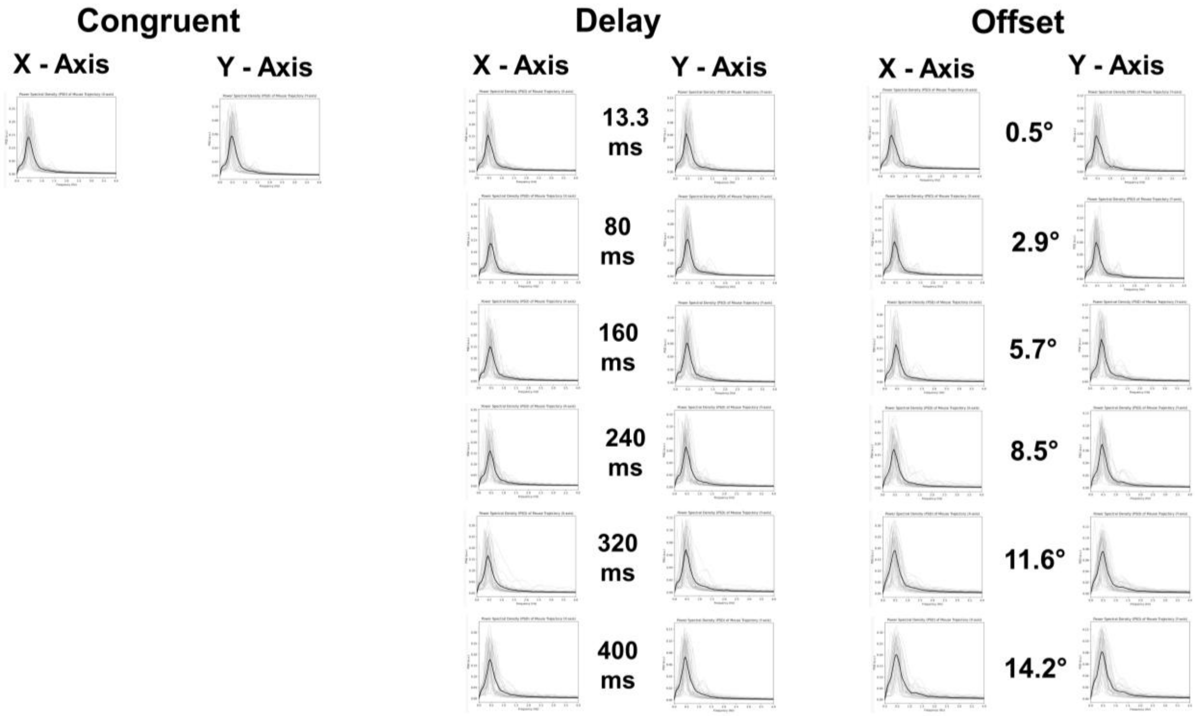

### Figure S3: Statistical comparison of power spectra

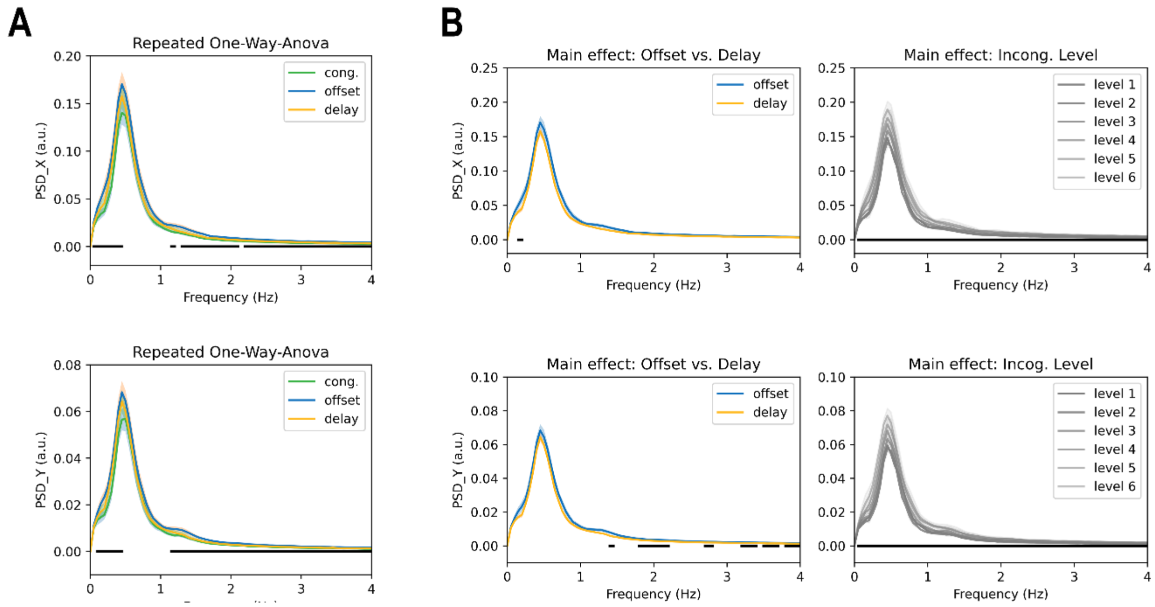


We used the MNE-python toolbox (Maris & Oostenveld, 2007; Gramford et al., 2013) to test for significant differences in the power spectral densities (PSDs) of executed movements between conditions. The PSDs were estimated with a frequency resolution of 0.05 Hz in the frequency range from 0 to 4 Hz. The cluster-forming threshold was set to $p < 0.001$, and the cluster-level significance threshold was set to $p < 0.05$. The plots show the averaged power spectra, with associated significances marked by black bars ($p<0.05$, corrected); separately for movements in the x-dimension (top rows) and y-dimension (bottom rows). A one-way repeated-measures ANOVA revealed significant differences across conditions (panel **A**). These were mainly driven by the congruent condition showing overall lower power; the power spectra of the delayed and offset conditions did not differ significantly around the key movement frequencies (i.e., 0-1 Hz), see the 2x6 repeated-measures ANOVA with the factors condition (offset, delay) and mismatch level (i.e.; the six respective levels of offset and delay) in **B**. There was a significant main effect of mismatch level, with stronger mismatches showing overall stronger power. We note that the present data were not very well suited to this kind of analysis; i.e., as shorter trials were likely to generate less consistent PSDs (and, therefore, overall lower power). Trial duration decreased linearly with mismatch level (see Results), which could explain the main effect of mismatch level shown above. In sum, for our study it is of primary importance that the PSDs showed similar shapes and an absence of higher-frequency components.

**References**

Eric Maris and Robert Oostenveld. Nonparametric statistical testing of EEG- and MEG-data. Journal of Neuroscience Methods, 164(1):177–190, 2007. doi:10.1016/j.jneumeth.2007.03.024.

Alexandre Gramfort, Martin Luessi, Eric Larson, Denis A. Engemann, Daniel Strohmeier, Christian Brodbeck, Roman Goj, Mainak Jas, Teon Brooks, Lauri Parkkonen, and Matti S. Hämäläinen. MEG and EEG data analysis with MNE-Python. Frontiers in Neuroscience, 7(267):1–13, 2013. doi:10.3389/fnins.2013.00267.

**Figure S4. Correlations between classification performances for delays (left) and offsets (right) in the execution vs observation phases**

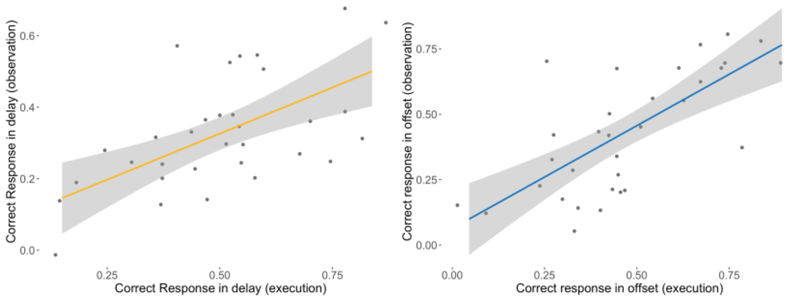


For delays (left), performance in the execution part significantly correlated with performance in the observation part (β = 0.54, *SE* = 0.12, *t* = 4.36, *p* < .001). For offsets (right), performance in the execution part significantly correlated with performance in the observation part (β = 0.81, *SE* = 0.15, *t* = 5.48, *p* < .001).

**Table S4.** Effect sizes relative to classification above chance level in the execution phase.

| | **Classification** | **Incongruence discrimination** |
|---|---|---|
| | *OR* | *Cohen's d* |
| | *Congruent* | |
| | 12.12 | - |
| | *Delay* | |
| 13.3 ms | 0.18 | -0.14 |
| 80 ms | 0.58 | 0.38 |
| 160 ms | 2.56 | 0.66 |
| 240 ms | 4.24 | 0.71 |
| 320 ms | 5.23 | 0.69 |
| 400 ms | 4.87 | 0.63 |
| Mean | 1.96 | 0.4 |
| | *Offset* | |
| 0.5° | 0.18 | -0.52 |
| 2.9° | 0.54 | -0.25 |
| 5.7° | 2.27 | 0.38 |
| 8.5° | 3.96 | 0.53 |
| 11.6° | 3.6 | 0.41 |
| 14.2° | 3.31 | 0.4 |
| Mean | 1.68 | 0.16 |

**Table S5, related to Table 1.** Comparison of delay vs offset classification and discrimination at each stimulus level in the execution phase, with associated standard errors of the mean in brackets.

| | **Delay** (Prob. correct) | **Offset** (Prob. correct) | ***p*** **delay vs offset** | ***effect size delay vs offset*** |
|---|---|---|---|---|
| | *Classification* | | | *Odds ratio* |
| 13.3 ms / 0.5° | 0.08 (0.02) | 0.08 (0.02) | >.99 | 0.99 |
| 80 ms / 2.9° | 0.23 (0.03) | 0.21 (0.03) | >.99 | 0.92 |
| 160 ms / 5.7° | 0.56 (0.04) | 0.53 (0.04) | >.99 | 0.89 |
| 240 ms / 8.5° | 0.68 (0.04) | 0.66 (0.04) | >.99 | 0.93 |
| 320 ms / 11.6° | 0.72 (0.04) | 0.64 (0.04) | .49 | 0.69 |
| 400 ms / 14.2° | 0.71 (0.04) | 0.62 (0.04) | .39 | 0.68 |
| Mean | 0.50 (0.03) | 0.46 (0.03) | .15 | 1.16 |
| | *Incongruence discrimination* | | | *Cohen's d* |
| 13.3 ms / 0.5° | 0.45 (0.08) | 0.32 (0.08) | .75 | 0.09 |
| 80 ms / 2.9° | 0.63 (0.07) | 0.41 (0.07) | .07 | 0.19 |
| 160 ms / 5.7° | 0.73 (0.07) | 0.64 (0.06) | .81 | 0.09 |
| 240 ms / 8.5° | 0.75 (0.06) | 0.69 (0.06) | .97 | 0.06 |
| 320 ms / 11.6° | 0.75 (0.06) | 0.65 (0.06) | .78 | 0.09 |
| 400 ms / 14.2° | 0.73 (0.06) | 0.64 (0.06) | .88 | 0.08 |
| Mean | 0.69 (0.04) | 0.58 (0.04) | **.002** | 0.24 |

Significant p-values are in bold font. All p-values are Holm-Bonferroni adjusted for multiple comparisons.

**Table S6.** Effect sizes relative to chance level in the observation phase and effect sizes of differences between execution and observation, related to Table 2.

| | Classification | Classification exec. vs obs. | Discrimination | Discrimination exec. vs obs. |
|---|---|---|---|---|
| | *OR* | *OR* | *Cohen's d* | *Cohen's d* |
| | | *Congruent* | | |
| | 2.88 | 0.24 | - | - |
| | | *Delay* | | |
| 13.3 ms | 0.82 | 4.57 | 0.73 | - 0.31 |
| 80 ms | 1.01 | 1.73 | 0.36 | 0.01 |
| 160 ms | 0.98 | 0.38 | 0.43 | 0.08 |
| 240 ms | 0.91 | 0.21 | 0.23 | 0.17 |
| 320 ms | 0.86 | 0.16 | 0.24 | 0.16 |
| 400 ms | 1.06 | 0.22 | 0.42 | 0.08 |
| Mean | 0.94 | 0.48 | 0.29 | 0.11 |
| | | *Offset* | | |
| 0.5° | 0.35 | 1.97 | - 0.43 | - 0.02 |
| 2.9° | 0.57 | 1.06 | - 0.28 | 0.01 |
| 5.7° | 1.43 | 0.63 | - 0.003 | 0.13 |
| 8.5° | 2.32 | 0.58 | 0.19 | 0.11 |
| 11.6° | 2.96 | 0.82 | 0.48 | - 0.02 |
| 14.2° | 0.94 | 0.94 | 0.38 | 0.01 |
| Mean | 1.45 | 0.86 | 0.04 | 0.13 |

### Control analysis with fixed trial lengths

In our comparison between execution and observation phases, we limited the trial duration during playback (to prevent participants from inferring conditions from differences in trial length). Here, we calculated a control analysis limited to only trials with a duration of <8s; i.e., excluding the other trials from the execution and the observation part. This analysis can be seen as excluding trials with very high uncertainty; i.e., where participants took very long to reach a decision. Note that this control analysis may suffer from reduced power due to the lower trail number; and that the number of excluded trials differed across the different mismatch levels ($\chi^2_{(12)}$=91.1, $p$<.001; but not across conditions, $\chi^2_{(2)}$=4.21, $p$=.12). These results should, therefore, be considered with some caution.

**Table S7, related to Table 1.** Mean rating performances in the execution phase, with associated standard errors of the mean in brackets. This table is provided only as a comparison for Table S8.

| | **Classification** | | **Incongruence discrimination** | |
|---|---|---|---|---|
| | Prob. correct | *p* > chance (33.3%) | Prop. correct | *p* > chance (50%) |
| | | *Congruent* | | |
| | 0.95 (0.01) | **<.0001** | - | - |
| | | *Delay* | | |
| 13.3 ms | 0.02 (0.02) | >.99 | 0.35 (0.15) | .84 |
| 80 ms | 0.21 (0.07) | >.99 | 0.74 (0.13) | .06 |
| 160 ms | 0.7 (0.07) | **<.0001** | 0.73 (0.09) | **.01** |
| 240 ms | 0.81 (0.05) | **<.0001** | 0.78 (0.09) | **.004** |
| 320 ms | 0.77 (0.06) | **<.0001** | 0.74 (0.09) | **.01** |
| 400 ms | 0.77 (0.05) | **<.0001** | 0.74 (0.08) | **.009** |
| Mean | 0.6 (0.05) | **<.0001** | 0.73 (0.06) | **<.001** |
| | | *Offset* | | |
| 0.5° | 0.05 (0.03) | >.99 | 0.56 (0.15) | .35 |
| 2.9° | 0.27 (0.07) | >.99 | 0.67 (0.11) | .13 |
| 5.7° | 0.61 (0.07) | **<.001** | 0.68 (0.09) | .11 |
| 8.5° | 0.63 (0.07) | **.0001** | 0.65 (0.08) | .13 |
| 11.6° | 0.69 (0.06) | **<.0001** | 0.64 (0.08) | .13 |
| 14.2° | 0.67 (0.06) | **<.0001** | 0.66 (0.08) | .13 |
| Mean | 0.54 (0.05) | **<.0001** | 0.66 (0.06) | **.004** |

Significant p-values are in bold font. All p-values are Holm-Bonferroni adjusted for multiple comparisons.

**Table S8, related to Table 2.** Mean rating performances in the observation phase, with associated standard errors of the mean in brackets, and comparisons against the execution phase.

| | Classification | | | Incongruence discrimination | | |
|---|---|---|---|---|---|---|
| | Prob. correct | *p* > chance (33.3%) | *p* execution vs observation | Prop. correct | *p* > chance (50%) | *p* execution vs observation |
| | | | *Congruent* | | | |
| | 0.66 (0.05) | **<.0001** | **<.0001** | - | - | - |
| | | | *Delay* | | | |
| 13.3 ms | 0.37 (0.07) | >.99 | **<.0001** | 0.71 (0.1) | .09 | **.03** |
| 80 ms | 0.31 (0.08) | >.99 | .30 | 0.7 (0.11) | .16 | .80 |
| 160 ms | 0.35 (0.07) | >.99 | **<.0001** | 0.6 (0.09) | .23 | .20 |
| 240 ms | 0.34 (0.07) | >.99 | **<.0001** | 0.65 (0.09) | .16 | .18 |
| 320 ms | 0.29 (0.06) | >.99 | **<.0001** | 0.58 (0.09) | .24 | .14 |
| 400 ms | 0.37 (0.07) | >.99 | **<.0001** | 0.71 (0.09) | .07 | .74 |
| Mean | 0.34 (0.05) | .38 | **<.0001** | 0.65 (0.06) | **.01** | .11 |
| | | | *Offset* | | | |
| 0.5° | 0.08 (0.04) | >.99 | .34 | 0.21 (0.1) | >.99 | **.04** |
| 2.9° | 0.2 (0.06) | >.99 | .38 | 0.38 (0.1) | >.99 | **.02** |
| 5.7° | 0.52 (0.08) | **.02** | .30 | 0.6 (0.08) | .32 | .44 |
| 8.5° | 0.64 (0.07) | **<.0001** | .87 | 0.65 (0.08) | .14 | .97 |
| 11.6° | 0.62 (0.06) | **<.0001** | .26 | 0.68 (0.08) | .08 | .65 |
| 14.2° | 0.66 (0.06) | **<.0001** | .78 | 0.67 (0.08) | .09 | .90 |
| Mean | 0.51 (0.05) | **<.001** | .31 | 0.57 (0.06) | .10 | .05 |

Significant p-values are in bold font. All p-values are Holm-Bonferroni adjusted for multiple comparisons.

**Table S9, related to Fig. 4B.** Results of the linear regression analysis on classification performance vs mismatch level in both phases (top) and the rmANOVA on the resulting regression weights (bottom).

| | Classification | | | | Incongruence discrimination | | | |
|---|---|---|---|---|---|---|---|---|
| | ß (*SE*) | *t- statistic* | *p-value* | $R^2$ | ß (*SE*) | *t- statistic* | *p-value* | $R^2$ |
| | | | | *Execution* | | | | |
| Delay | 0.72 (0.09) | 7.99 | **<.001** | 0.32 | -0.02 (0.12) | -0.17 | .87 | 0.0003 |
| Offsett | 0.53 (0.1) | 5.41 | **<.001** | 0.42 | -0.04 (0.14) | -0.32 | .75 | 0.0008 |
| | | | | *Observation* | | | | |
| Delay | 0.07 (0.08) | 0.79 | .43 | 0.005 | -0.05 (0.1) | -0.52 | .6 | 0.003 |
| Offset | 0.52 (0.09) | 5.49 | **<.001** | 0.16 | 0.26 (0.011) | 2.28 | **.02** | 0.04 |

**Figure S5, related to Fig. 4B.** Results of the rmANOVA on the regression weights resulting from the above (Table S8).

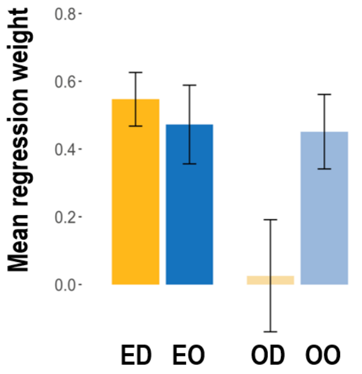


A rmANOVA with the factors experimental phase (execution, observation) and feedback condition (delay, offset) on the resulting regression weights replicated the significant main effect ($F(1,27)=5.27$, $p=.03$ $\eta_p^2 = 0.16$), with steeper slopes in the execution than in the observation phase; and the significant interaction ($F(1,27)=5.32$, $p=.03$, $\eta_p^2 = 0.17$).
